# CX3CL1 and CX3CR1 Expressing Tendon Cells – A novel Immune Cell Population in the Tendon Core

**DOI:** 10.1101/693317

**Authors:** Christine Lehner, Gabriel Spitzer, Renate Gehwolf, Andrea Wagner, Nadja Weissenbacher, Christian Deininger, Katja Emmanuel, Florian Wichlas, Herbert Tempfer, Andreas Traweger

## Abstract

Tendon disorders frequently occur and recent evidence has clearly implicated the presence of immune cells and inflammatory events during early tendinopathy. However, the origin and properties of these cells remain poorly defined. Therefore, the aim of this study was to determine the presence of myleoid cells in healthy rodent and human tendon tissue and to characterize them. Using various transgenic reporter mouse models, we demonstrate the presence of tendon cells in the dense matrix of the tendon core expressing the fractalkine (Fkn) receptor CX3CR1 and its cognate ligand CX3CL1/Fkn. Pro-inflammatory stimulation of 3D tendon-like constructs *in vitro* resulted in a significant increase in the expression of IL-1β, IL-6, Mmp3, Mmp9, Cx3cl1, and epiregulin which has been reported to contribute to inflammation, wound healing, and tissue repair. Furthermore, we demonstrate that inhibition of the fractalkine receptor blocked tendon cell migration *in vitro* and show the presence of CX3CR1/CX3CL1/EREG expressing cells in healthy human tendons. Taken together, we demonstrate the presence of CX3CL1+/CX3CR1+ “tenophages” within the healthy tendon proper potentially fulfilling surveillance functions in tendons.

**Summary Statement:** Here, we demonstrate the presence of a macrophage-like, CX3CL1/CX3CR1-expressing cell population within the healthy tendon proper potentially fulfilling a surveillance function.

## INTRODUCTION

Tendon pathologies and injuries are one of the most common musculoskeletal disorders, however due to the tissue’s poor regenerative capacity the healing process is long-lasting and outcomes are often not satisfactory. Consequently, tendinopathies represent a substantial social and economic burden (Schneider et al., 2018). The limited availabilty of effective treatment options not only ows to the multifactorial nature of tendinopathies, but above all results from our insufficient understanding of the cellular and molecular mechanisms leading to the onset and progression of the disease. Therefore, gaining a deeper insight into the nature and function of tendon-resident cells in tissue homeostasis and disease is imperative for developing new treatment strategies for tendinopathies.

Due to the composition and structure of the extracellular matrix (ECM), tendons are able to withstand enormous tensile forces, so that spontaneous ruptures rarely occur without preceding features of tissue degeneration. Besides repetitive overload, smoking, and the intake of certain drugs, also obesity and various metabolic diseases are recognized risk factors for the development of tendinopathies. Interestingly, a role of inflammation in the pathogenesis of tendinopathy has long been debated, the underlying mechanisms being poorly understood. The presence of myeloid and lymphoid cells such as mast cells, T cells, and macrophages during early human tendinopathy however highlight a role of inflammation in tendon disease (Dean et al., 2016; Kragsnaes et al., 2014; Millar et al., 2010). However, the origin of these immune cells is unclear; whether they invade the tissue from the circulation and neighbouring tissue, or whether tissue-resident myeloid cells are present and are activated upon damage, or a combination of both mechanisms. Generally, tissue-resident macrophages *in vivo* are not a homogeneous cell population, but heterogeneous in nature and respond to certain stimuli with overlapping functions and phenotypes and therefore often can not be classified into simple, polarized categories (Davies and Taylor, 2015). As the majority of these cells are usually situated in the vicinity of blood vessels (Hume et al., 1984), it seems plausible that this would also apply for tissue-resident myeloid cells in tendons. However, the presence and distribution of immune cells in healthy tendons has not been thoroughly investigated so far and due to the hypovascular nature of tendons, we hypothesize that in tendons resident myeloid cells not only are present in the perivascular region, but also reside within the dense, collagen-rich tendon core fulfilling a surveillance function similar to Langerhans cells in the skin or microglia in the brain (Deckers et al., 2018; Lehner et al., 2016).

In general, the main effectors of inflammation are myeloid cells, most notably monocytes and macrophages. Among the known factors that control e.g. monocyte recruitment is the chemokine CX3CL1, or Fractalkine (FKN), and its cognate receptor CX3CR1 (Lee et al., 2018). CX3CR1 is expressed by myeloid and lymphoid lineage cells, including mast cells and natural killer cells (Mass et al., 2016; Sasmono and Williams, 2012). In addition, CX3CL1/FKN has been demonstrated to regulate the communication between neurons, glia and microglia, and CX3CR1-expressing microglia have been suggested to be pivotal in limiting tissue injury during inflammation and neuro-degeneration (Sheridan and Murphy, 2013). Overall, depending on the tissue type CX3CR1-expressing cells can either contribute to maintenance of tissue homeostasis or play a role in disease progression. These findings prompted us to investigate if the CX3CL1/CX3CR1 axis might also be relevant in tendons. Therefore, the purpose of this study was (1) to assess the presence of tendon core-resident cells in healthy rodent and human tissues expressing immune cell-related markers and (2) to explore the ramifications of pro-inflammatory stimulation on the CX3CL1/CX3CR1 system in 3D tendon-like constructs *in vitro.*

## RESULTS

### Tendon-resident cells express immune cell-related markers

To evaluate the presence of tendon-resident cells expressing immune-cell markers we probed Achilles tendon tissue sections from the transgenic *Scx-GFP* tendon reporter mouse strain (Pryce et al., 2007). As shown in figure 1A and B, GFP-positive cells located in the dense tendon core co-expressed the widely used pan-macrophage marker CD68, F4/80, a unique marker of murine macrophages, and also the macrophage-specific hemoglobin (Hb) scavenger receptor CD163. Further, immunohistochemical staining also revealed tendon cells co-expressig MHC class II, a membrane-bound marker for antigen-presenting cells such as macrophages, B-lymphocytes and dendritic cells (Kristiansen et al., 2001). To further substantiate the presence of mlyeoid cells in the tendon proper we also investigated Achilles tendon tissue of the transgenic MacGreen reporter mouse strain. These mice express EGFP under the control of the mouse colony stimulating factor 1 receptor (*Csf-1r*) promoter, labelling mononuclear phagocyte lineage cells (Sasmono et al., 2003). As shown in figure **1C** several cells in the tendon proper were positive for EGFP, indicating the presence of myeloid cells. Further, the majority of the EGFP-positive cells also stained positive for Cx3cr1 (Fkn receptor) and expression of the receptor was also confirmed using a transgenic mouse strain expressing EGFP driven by the *Cx3cr1* promoter (Jung et al., 2000) (**Suppl. fig. 1**). Finally, by employing double immunolabelling we further demonstrate that the fractalkine receptor and its ligand Cx3cl1 are both co-expressed by tendon cells (**Fig. 2A**) and the expression of Cx3cr1 specifically in tendon cells was also confirmed by probing Achilles tendon sections of the *Scx-GFP* tendon reporter mouse strain (**Fig. 2B**).

**Fig. 1:**
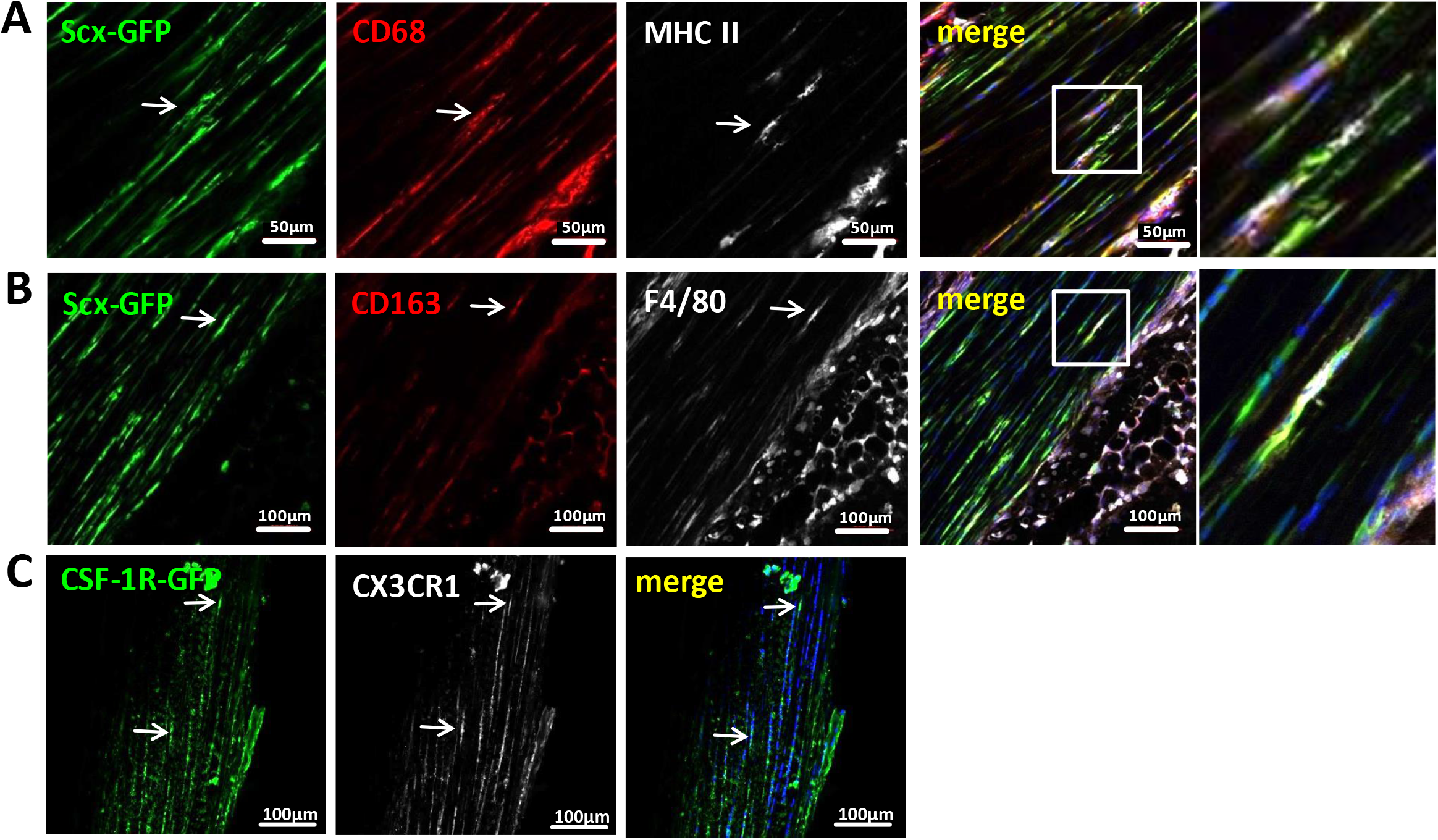
Immunohistochemical staining of immune cell markers on histological sections of Achilles tendons from *Scleraxis (SCX)-GFP* transgenic mice reveals that SCX-positive cells co-express CD68, MHCII, CD163, and F4/80, respectively (A, B; arrows). Cryo-sections of Achilles tendon from transgenic *Csf-1r* and *Cx3cr1-GFP* reporter mice show that cells within the dense part of the tendon are positive for CSF-1R and CX3CR1 (arrows).

**Fig. 2:**
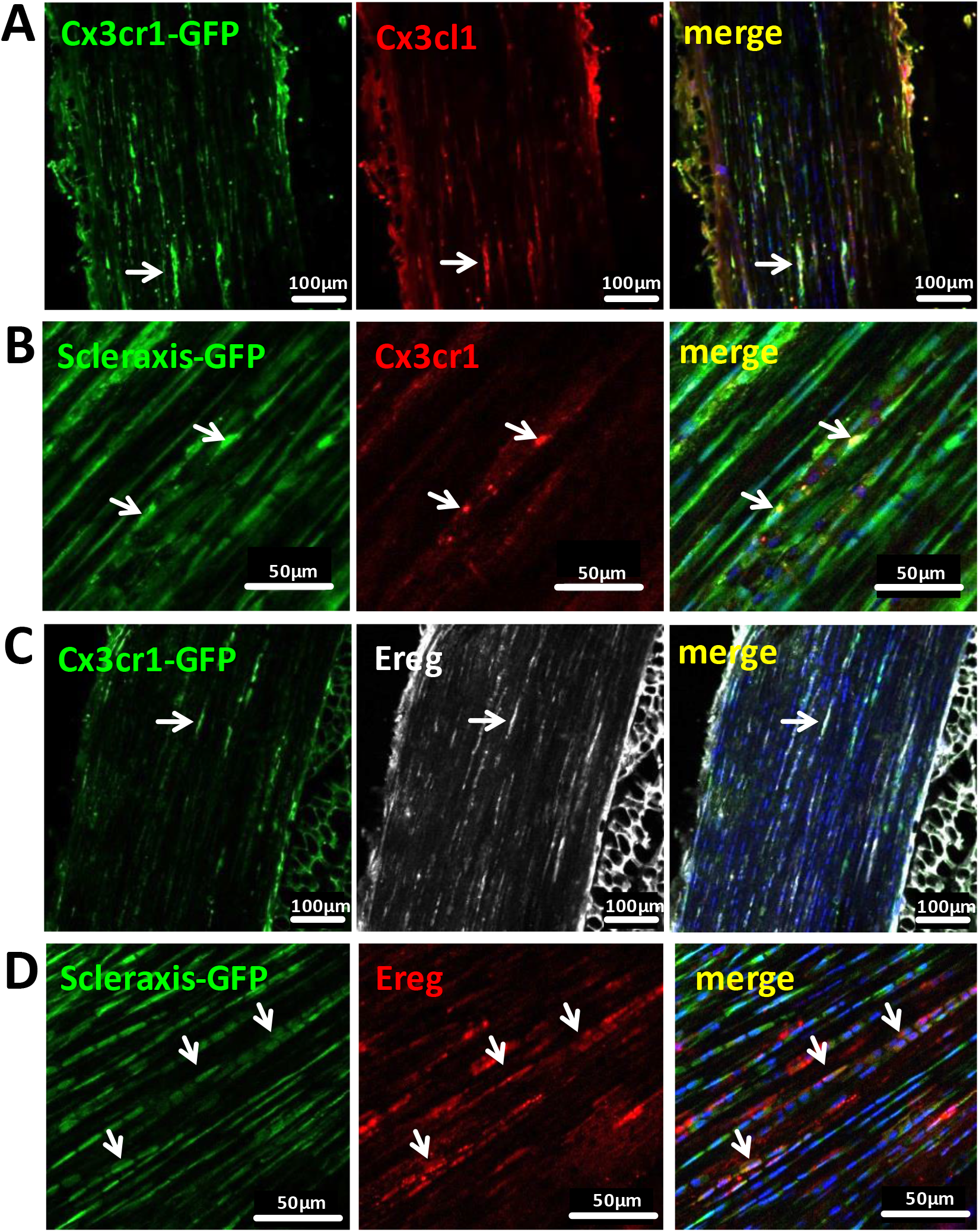
Cryosections of Achilles tendons from transgenic *Cx3cr1-GFP* (A, C) and *Scx-GFP* (B, D) reporter mice immunohistochemically stained with antibodies recognizing CX3CL1/FKN, its receptor CX3CR1, and EREG respectively. Arrows point towards cells co-expressing the respective proteins.

FKN has been described to induce shedding of epiregulin (EREG), a 46-amino acid protein belonging to the Epidermal Growth Factor (EGF) family of peptide hormones, and further to rapidly increase epiregulin mRNA expression 20-fold (White et al., 2010). Therefore, we investigated tendon tissue sections for the presence of EREG. Indeed, epiregulin is also expressed in tendon-resident cells expressing Scx-GFP or Cx3cr1-EGFP (**Fig. 2C, D**). Finally, Cx3cr1-positive cells also express both macrophage markers CD68 and CD163 (**Suppl. fig. 2**).

Next, to determine whether these cells, apart from their macrophage-associated marker profile, also possess phagocytic activity we exposed unfixed rat flexor tendons to pHrodo™ Green S. aureus Bioparticles™ which upon cellular uptake emit fluoresecence due to a shift in pH. As shown in **figure 3**, we detected several positive cells within the tendon core embedded in the dense collagenaous matrix, demonstrating the presence of phagocytic cells within the tendon proper *in vivo.*

**Fig. 3:**
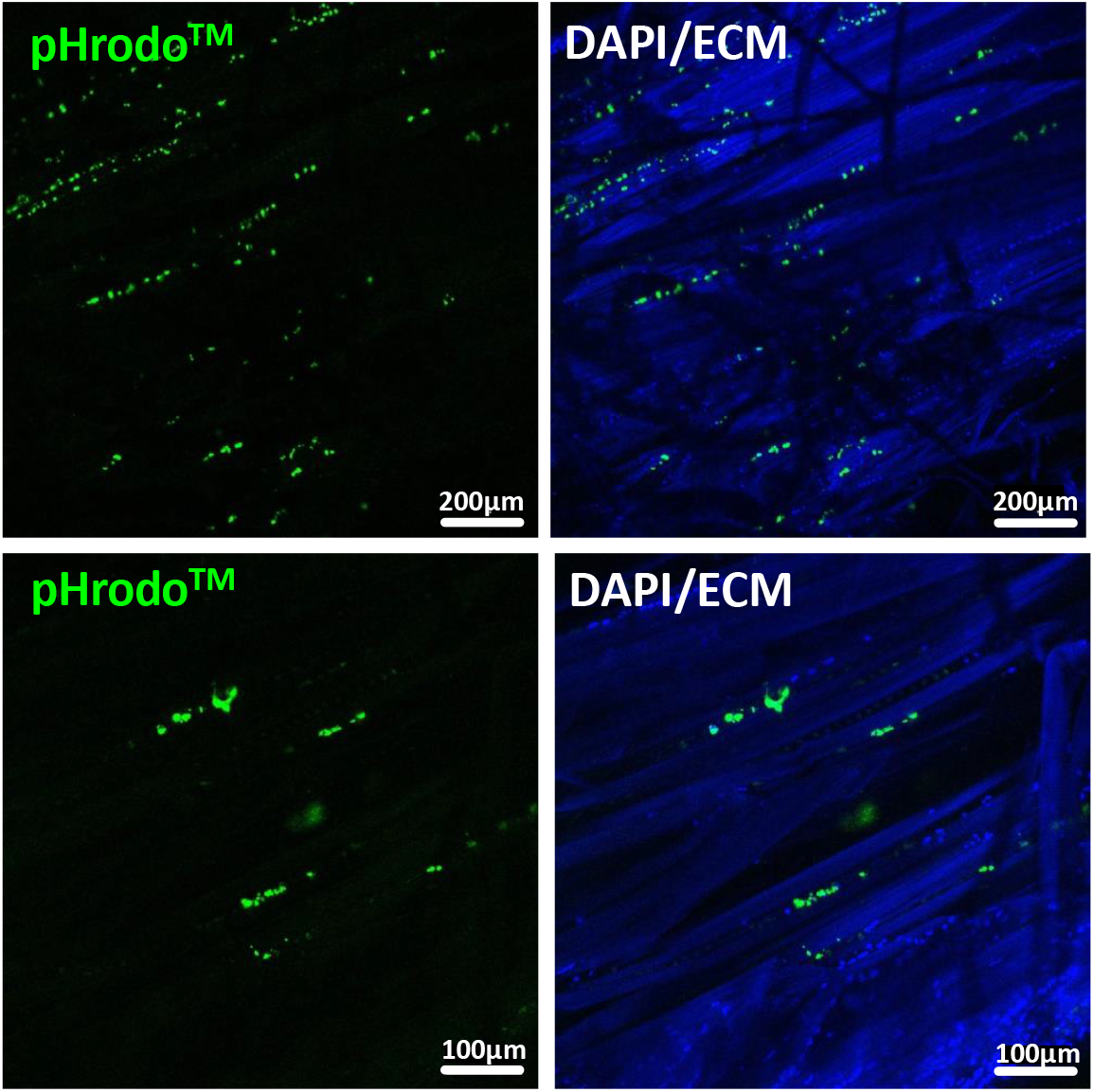
In situ phagocytosis assay on unfixed rat flexor tendons shows that tendon cells lying within the dense collagen matrix (shown by extracellular matrix (ECM) autofluorescence / blue channel) exert phagocytic activity (green fluorescence). Two representative regions are shown.

### Pro-inflammatory stimulation of 3D tendon-like constructs increases fractalkine and epiregulin expression

Having identified tendon-resident cells expressing immune cell-related markers, we next examined the response of primary tendon stem and progenitor cells (subsequently referred to as TDSPCs) to pro-inflammatory stimuli. We therefore generated 3D type I collagen-embedded tendon cell cultures as previously described (Gehwolf et al., 2019) and analyzed the expression of both tendon-specific and matrix-associated as well as inflammation-related markers after exposure to IL-1β, TNF-α or a combination of both (**Fig. 4A**). As shown in figure **4B**, stimulation of the constructs significantly increased the gene expression of *IL-1β, TNF-α*, and *IL-6*, as well as several extracellular-matrix (ECM)-associated proteins such as lysyloxidase (Lox) and the matrix metalloproteinases (MMPs) *Mmp1, Mmp3,* and *Mmp9*. A synergistic effect of Il-1β and TNFα stimulation was seen for several candidate genes, however IL-1β-treatment generally had a more pronounced effect on gene expression. No significant effect was evident for the expression of type I collagen (*Col1a1*) and type 3 collagen (*Col3a1*). Further, there was little or no impact on the expression of the tenogenic marker proteins Tenomodulin (*Tnmd*), Mohawk (*Mkx*) and Scleraxis (Scx).

**Fig. 4:**
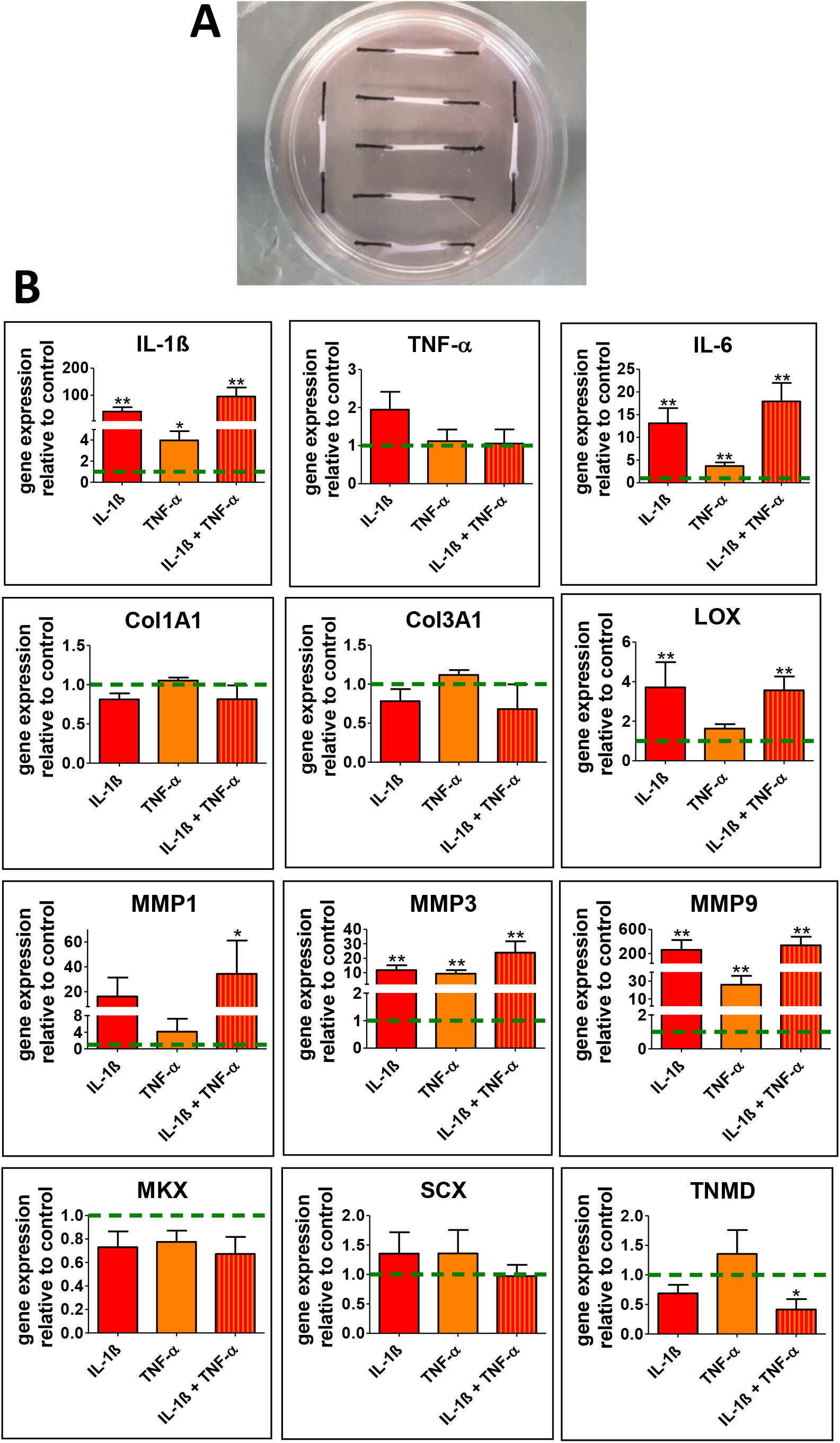
3D tendon-like constructs were stimulated with either IL-1β, TNF-α, or a combination of both cytokines (A). Effects on the expression levels of genes encoding for inflammatory proteins (*IL-1β, TNFa, IL-6*), extracellular matrix-related proteins (e.g. *Col1a1, Col3a1, Lox, Mmp-1, Mmp-3, Mmp-9*), and tendon cell-related marker proteins (*Mwk, Scx, Tnmd*) were assessed by qPCR. Significant changes were detected for *IL-1β, IL-6, Lox, Mmp-1, Mmp-3, and Mmp-9.* Bars represent mean ± SEM (for 5 individual animals); *p<0.05, **p<0.01, Mann Whitney test. Dashed green line: control reference.

IL-1β exposure led to a moderate 2-fold increase in the expression of the macrophage-related marker CD68, whereas a significant increase (≥20-fold) in *Fkn* (*Cx3cl1*) and *Ereg* mRNA quantitites was observed, which was even higher if co-stimulated with TNFα. These results were further underscored by immunofluorescent analysis, demonstrating that pro-inflammatory treatment mainly affected the expression of Cx3cl1 and Ereg (**Fig. 5B**). Finally, to obtain quantitative data on protein levels we also performed Western blot analysis on lysates prepared from stimulated and unstimulated 3D tendon-like constructs. Again, a significant increase in expression was observed for both Cx3cl1 and Ereg (**Fig. 5C**).

**Fig. 5:**
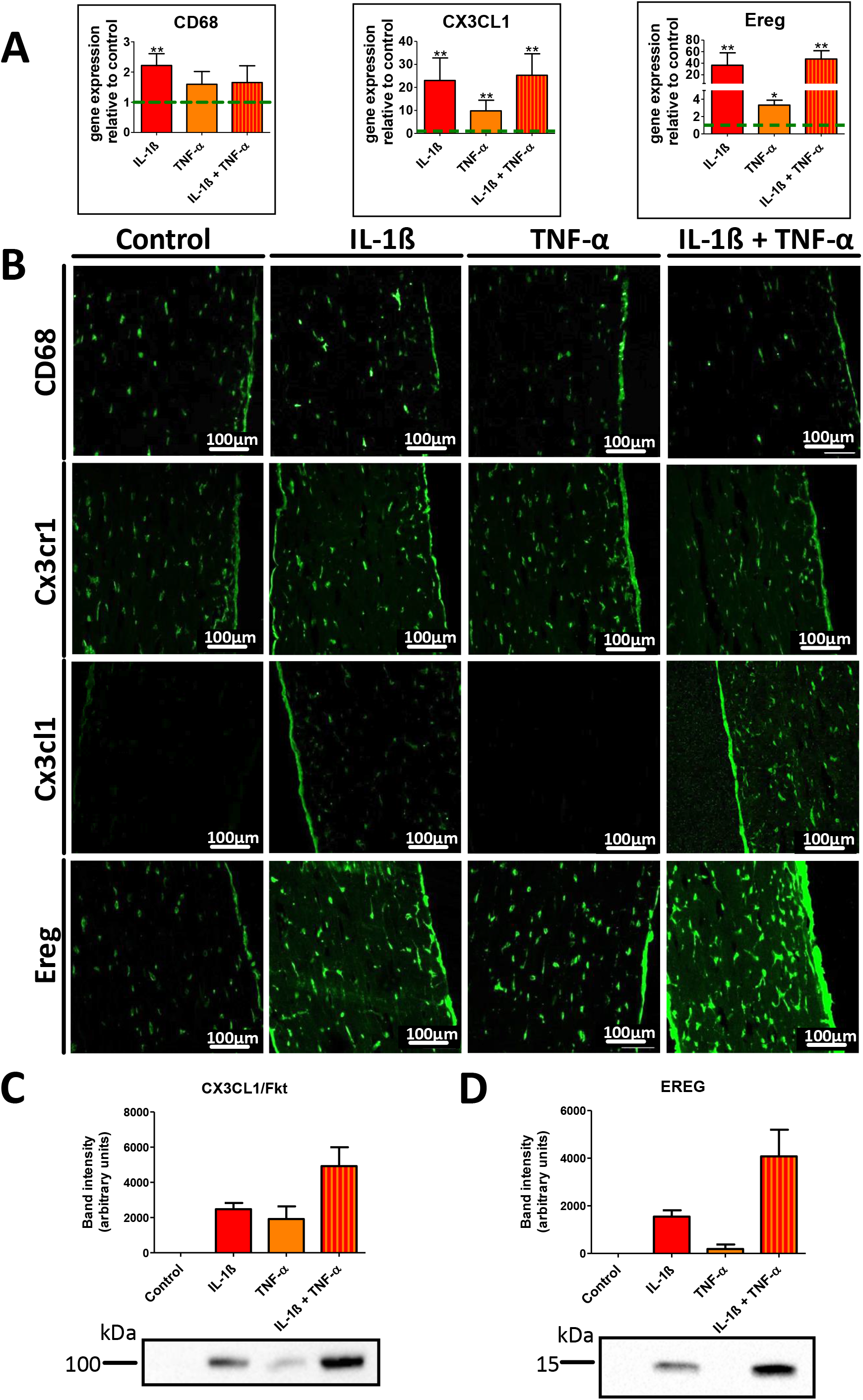
Effects of pro-inflammatory stimulation of tendon-like constructs on mRNA (A), and protein level (B, C). IL-1β or TNF-α or a combination of both cytokines resulted in a significant upregulation of *CD68, Cx3cl1,* and *Ereg* mRNA expression (A). Immunohistochemical stainings confirmed the qPCR findings. Cx3cr1 remained unaffected by the treatment (B). Furthermore, Western blot analysis revealed a synergistic effect of IL-1β and TNF-α on Cx3cl1 and Ereg expression (C). *p<0.05, **p<0.01, Mann-Whitney test.

### Inhibition of CX3CR1 signalling blocks tendon cell migration

In order to address a putative function of the CX3CL1/CXCR1 signalling axis in tendon-resident cells, we next performed cell migration assays. To inhibit CX3CR1 we applied AZD 8797 (Axon Medchem, Groningen, Netherlands), a selective, high-affinity small-molecule inhibitor of CX3CR1. Importantly, early passage rat TDSPCs (p1) retain the expression of both FKN and its receptor (**Fig. 6A**). Interestingly, treatment with the FKN receptor antagonist led to a reduction of IL-1β-triggered mRNA expression of *IL-1β* and *IL-6* back to control levels (**Fig. 6B/C**). Analysis of the wound scratch assay revealed that AZD 8797 almost completely blocked migration of TDSPCs on uncoated and type I collagen-coated cell culture dishes (**Fig. 6D/E**).

**Fig. 6:**
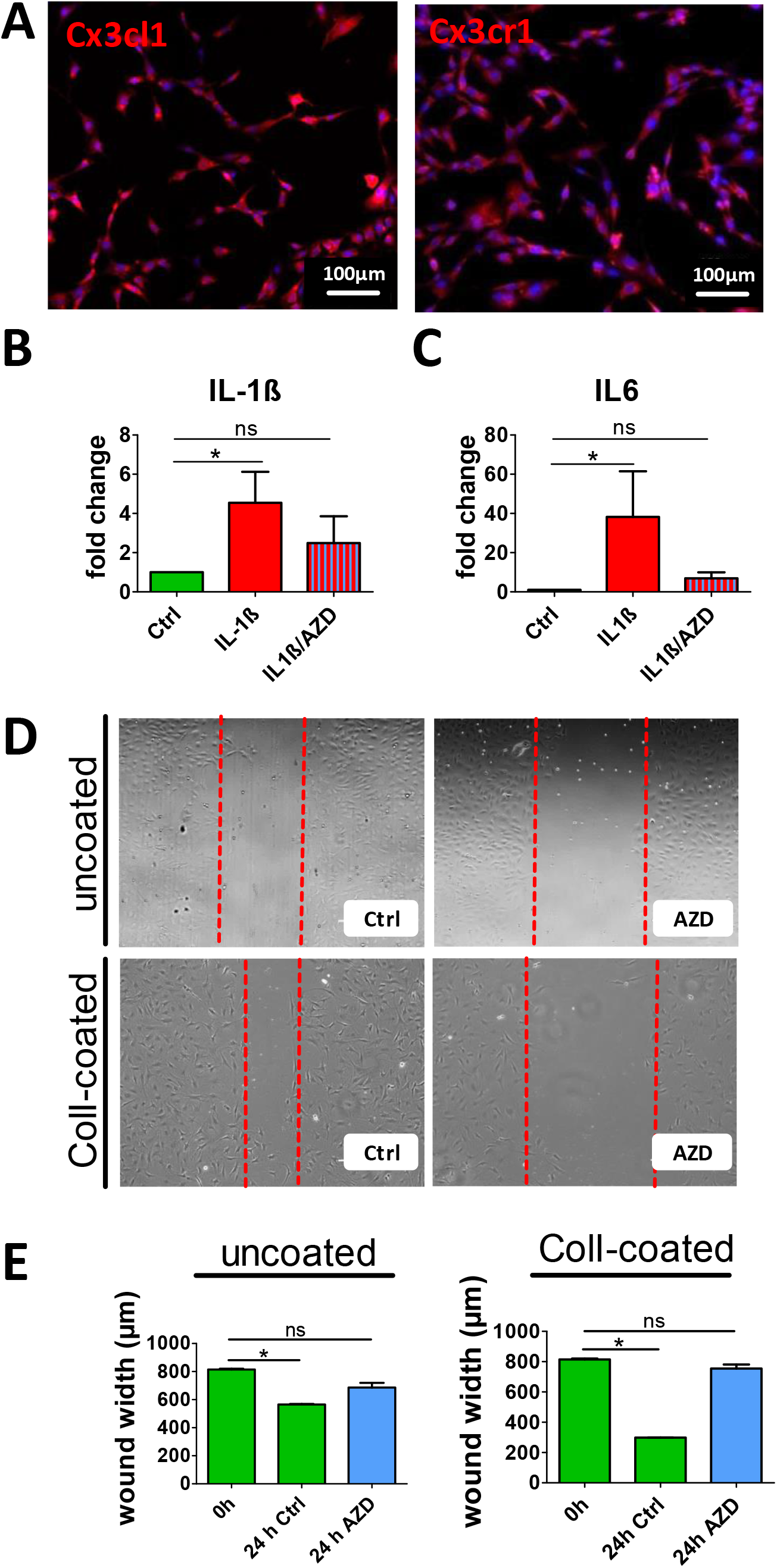
Rat tendon-derived cells (passage 1) express Cx3cl1 and Cx3cr1 (A). Addition of AZD 8797 attenuates IL-1β triggered upregulation of both IL-1β and IL-6 (B, C). Representative images (D) showing wound scratch assays on uncoated and collagen-coated culture plates. Quantitative analysis revealed that the FKN inhibitor AZD 8797 significantly reduces migration (E). *p<0.05, **p<0.01, Kruskal Wallis and Dunn’s Multiple Comparison test.

### CX3CL1, CX3CR1, and epiregulin are expressed in healthy human tendon tissue

Finally, we were interested to see whether fractalkine, its receptor CX3CR1 and epiregulin are also expressed in healthy human tendons. To this end, we probed cryo-sections of human semitendinosus tendons obtained from a healthy, 34 year old male (**Fig. 7A**). Indeed, next to a strong expression at blood vessel walls (**Suppl. fig. 3**), our analysis revealed the presence of dinstinct cells within the tendon proper expressing CX3CL1, CX3CL1, and EREG (**Fig. 7B-D**). To conclude, our results clearly demonstrate the presence of a CX3CL1/CX3CR1/EREG expressing cell population in healthy murine and human tendon tissue.

**Fig. 7:**
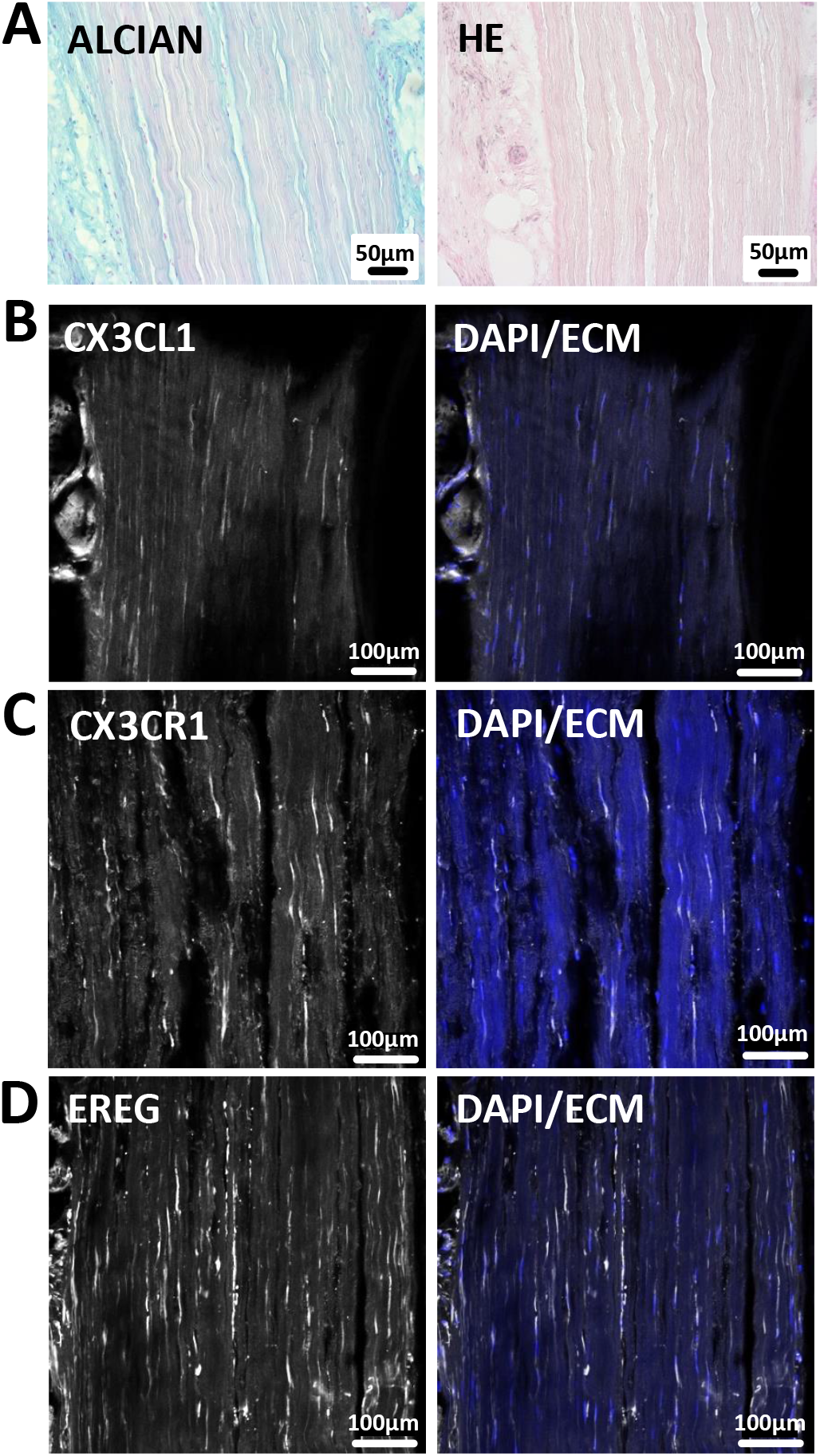
Cryo-sections of intact human semitendinosus tendon (♂, 34 years). Alcian Blue and Hematoxylin-Eosin (HE) stainings show the parallel alignment of collagen fibers and elongated cell nuclei characteristic for intact tendons (A). Immunofluorescent images demonstrating the presence of cells expressing CX3CL1/FKN (B), its cognate receptor CX3CR1 (C), and epiregulin (EREG) (D) in the tendon proper.

## DISCUSSION

Our understanding of the cellular and molecular mechanisms underlying tendinopathies remains very fragmentary. The term tendinopathy encompasses a broad sprectrum of tendon-related diseases and is mainly characterized by activity-related pain. Historically, there has been substantial debate about the terminology and if inflammation is of importance in the development and progression of tendinopathies (Khan et al., 2002; Khan et al., 2000). In contrast, more recent studies elegantly highlight the involvement of immune cells and activation of inflammatory processes in tendinopathy (Dean et al., 2017). However, the origin of these cells remains unknown and it is unclear if they mainly extravasate into the tissue upon injury or metabolic stress or if tendon-resident macrophages or mast cells exist in healthy tendon tissue initiating the first line response. Therefore, we aimed to formally demonstrate the presence of myleoid cells in intact, healthy murine and human tendons and to characterize a population of “tenophages” in the tendon core.

In the present study, we demonstrate the presence of cells positive for myleoid cell-related markers located within the dense tendon core region (**Fig. 1**). Interestingly, these cells were also positive for the widely accepted tendon-specific marker Scleraxis, a member of the basic helix-loop-helix (bHLH) superfamily of transcription factors. Hence, it is indeed tendon cells themselves expressing myeloid surface marker proteins. To our knowledge, this is the first description of such a cell population in healthy tendons. Apart from their cell surface marker profile these cells also appear to exert phagocytic activity as evidenced by an *ex vivo* phagocytosis assay (**Fig.3**). In addition, by making use of a transgenic mouse model we demonstrate the presence of a population of cells within the tendon proper, expressing Fkn (Cx3cl1) and its cognate receptor Cx3cr1 (**Fig.2**). CX3CR1 expression is associated with increased migration and site specific dissemination having been shown to be expressed by endothelial cells, mast cells, monocytes, tissue-resident macrophages, natural killer (NK) cells, microglial cells, neurons and subpopulations of T-lymphocytes (Imai et al., 1997; Papadopoulos et al., 2000; You et al., 2007). The seven-transmembrane domain G protein-coupled fractalkine receptor CX3CR1 mediates several intracellular signalling pathways, such as the p38MAPK signalling and the Akt pathway (Li et al., 2016; Wu et al., 2016). It has two known functional ligands, the chemokine CX3CL1 (also called neurotactin or fractalkine/FKN) and CCL26 (eotaxin-3), the latter being 10-fold less potent than CX3CL1 (Nakayama et al., 2010). FKN is structurally unique amongst the family of chemokines and is expressed both in the central nervous system and peripheral nerves, as well as in endothelial cells, dendritic cells and lymphocytes (Bazan et al., 1997; Kanazawa et al., 1999; You et al., 2007). It is constitutively cleaved by the ADAM-metalloprotease ADAM10 and upon cell stress, such as tissue injury, shedding is further promoted by ADAM17 (also known as the TNF-α converting enzyme, TACE), releasing an extracellular soluble fragment. In addition, the cysteine protease Cathepsin S has been shown to selectively cleave FKN (Fonovic et al., 2013). In its soluble form FKN mediates chemotaxis of immune cells, whilst membrane bound FKN acts as an adhesion molecule mediating leukocyte capture and infiltration (Clark et al., 2011; Imai et al., 1997; Umehara et al., 2004). FKN has been reported to be released by apoptotic lymphocytes stimulating macrophage chemotaxis and recruiting professional phagocytes to the site of cell death (Truman et al., 2008). Beyond simple recruitment, FKN has also been shown to enhance the ability of macrophages and microglia to execute their phagocytic functions (Tsai et al., 2014). Since accumulation of microruptures preceding tendon tears goes along with cell death and subsequent clearance of the cellular debris is required, it is tempting to speculate that the presence of FKN in the tendon might serve as "find-me” signal for macrophages invading the tissue from the circulation (Lundgreen et al., 2011; Sokolowski et al., 2014).

Geissmann et al. describe two different circulating monocyte populations, CCR2^-^CX_3_CR1^high^ monocytes (MCs) that home constitutively to tissues and short-lived CCR2^+^CX_3_CR1^low^ monocytes that only home to inflamed tissues (Geissmann et al., 2003). The authors suggest that cells derived from resident CCR2^-^CX_3_CR1^high^ monocytes, such as osteoclasts, Kupffer cells, and microglia, are involved in tissue homeostasis (Geissmann et al., 2003). Along these lines, CCR2^-^CX_3_CR1^high^ (also termed “nonclassical”) monocytes exhibit a unique ability to patrol the resting vasculature and remove debris (Auffray et al., 2007; Carlin et al., 2013). CX3CR1 positive cells in the tendon core might therefore represent such a population of myeloid precursor cells and indicate a role of CX3CR1 in tendon tissue homeostasis. It is also noteworthy, that non-classical MCs generally possess inflammatory characteristics and secrete inflammatory cytokines upon stimulation (Yang et al., 2014), similar to what we have observed for tendon cell-derived 3D constructs *in vitro* (**Fig. 4B**).

CCR2^-^CX_3_CR1^high^ monocytes have been described within the parenchyma of multiple tissues, including the brain. Sheridan and Murphy highlight the crosstalk of neurons and glia in health and disease and discuss that the FKN/CX3CR1 ligand/receptor pair seems to have evolved as a communication link between neurons and microglial cells, being crucial not only for maintaining tissue homeostasis under normal physiological conditions, but also being activated under inflammatory conditions such as stroke or Alzheimer’s disease (Sheridan and Murphy, 2013). We speculate that the observed presence of the CX3CL1/CX3CR1 system within the tendon might serve similar surveilling funtions as in the brain and that upon inflammatory stimulation the system reacts by upregulating FKN thereby attracting additional monocytes from the circulation.

Fractalkine has also been shown to induce aortic smooth muscle cell proliferation through an autocrine pathway by initiating phosphorylation of the mitogen-activated protein (MAP) kinases p38, c-Jun N-terminal kinase (JNK) and extracellular-regulated kinase (ERK) 1/2, as well as the serine-threonine kinase Akt in osteoarthritis fibroblasts (Klosowska et al., 2009; White et al., 2010). Interestingly, the observed effects of FKN on proliferation of coronary artery smooth muscle cells (CASMCs) are accompanied by transcription and release of epiregulin. In their study, White et al. describe that FKN induces shedding of epiregulin and increases epiregulin mRNA expression 20-fold within 2 hours (White et al., 2010). Here we report the presence of Scx-positive tendon cells also expressing Cx3cr1 and Ereg. Epiregulin is a 46-amino acid protein belonging to the Epidermal Growth Factor (EGF) family of peptide hormones. It binds to EGF receptors (EGFR) ErbB1 (HER1) and ErbB4 (HER4) and can stimulate signaling of ErbB2 (HER2/Neu) and ErbB3 (HER3) through ligand-induced heterodimerization with a cognate receptor. EREG is initially expressed as an extracellular transmembrane protein, which is cleaved by disintegrins and metalloproteinase enzymes (ADAMs) releasing a soluble form. Epiregulin has been shown to contribute to inflammation, wound healing, tissue repair, and oocyte maturation by regulating angiogenesis and vascular remodeling and by stimulating cell proliferation (Harada et al., 2015; Martin et al., 2017; Murakami et al., 2013; Riese and Cullum, 2014). In renal proximal tubular cells (RPTC), addition of 10ng/ml epiregulin enhanced both RPTC proliferation and migration via activation of the EGF receptor (EGFR), Akt, a downstream kinase of phosphoinositide 3-kinase (PI3K), and extracellular signaling-regulated kinase 1/2 (ERK1/2)(Zhuang et al., 2007). Similarly, for adipose derived mesenchymal stem cells epiregulin has been described to promote migration and chemotaxis ability via mitogen-activated protein kinase signalling pathways (Cao et al., 2018). Further, in Caco-2 epithelial cells EREG mRNA and protein levels have been shown to be increased by incubation with exogenous IL-1β (Massip-Copiz et al., 2018). This finding is well in line with our own data revealing that stimulation of 3D tendon-like constructs with IL-1β, or a combination of IL-1β and TNF-α significantly increased the expression of Ereg both on the gene as well as on the protein level.

Next to enhancing cell proliferation, FKN also promotes migration. Klosowska et al. demonstrate that FKN effectively induces migration of osteoarthritis (OA) fibroblasts (Klosowska et al., 2009). Similar findings have been reported by You et al. who showed that FKN is an angiogenic mediator *in vitro* and *in vivo.* FKN significantly induced migration of human umbilical vein endothelial cells (HUVECs) as well as bovine retinal capillary endothelial cells (BRECs) and promoted formation of endothelial cell capillary tubes on synthetic matrix. Moreover, FKN promoted blood vessel growth in a rabbit corneal pocket neovascularization assay (You et al., 2007). These observations of the pro-migratory effect of FKN corroborate our own data showing that addition of the CX3CR1 specific inhibitor AZD 8797 results in significantly reduced migration of rat tendon-derived cells in vitro (**Fig. 6**).

Interestingly, nuclear factor kappaB (NF-ĸB) signaling has recenty been demonstrated to be increased in clinical tendinopathy (Abraham et al., 2019) and it is known that FKN is stimulated by NF-ĸB-mediated inflammatory processes. Garcia et al. showed that NF-ĸB-dependent FKN induction in rat aortic endothelial cells is stimulated by IL-1β, TNF-α and lipopolysaccharide (LPS) (Garcia et al., 2000). Moreover, in human lung fibroblasts a dramatic increase in both soluble CX3CL1 protein and mRNA transcripts in a dose- and time-dependent manner has been reported to be synergistically induced by a combination of IL-1β and IFN-γ (Isozaki et al., 2011). Again, we observed similar responses in 3D tendon-cell cultures upon stimulation with IL-1β, TNF-α, or a combination of both (**Fig. 5**).

In summary, we describe the presence of macrophage-like tendon cells and provide evidence for the expression of the CX3CL1/CX3CR1 axis and the peptide hormone epiregulin in healthy rodent as well as human tendons. Interestingly, not only did we observe perivascular expression of these proteins, but also very distinctly in cells within the dense, collagen-rich matrix of tendons. We therefore propose that this newly identified cell population fulfils a surveillace function and is activated upon tendon tissue injury or pathological stress. Given the role in cell proliferation and angiogenesis upon inflammation and considering that both are hallmarks of tendinopathy, targeting the CX3CR1/CX3CL1/EREG axis could potentially open up new vistas in tendinopathy therapy.

### Materials and Methods

#### Cell culture

Primary TDSPCs were isolated from Achilles tendons of 5 rats (Fisher, female, 12 weeks). To this end, rat Achilles tendons were dissected, finely minced and digested in Dulbecco’s modified Eagle’s medium (DMEM) containing 2 mg/ml type II collagenase (Sigma-Aldrich, St. Louis, MO, USA) for 12 hours at 37 °C and 5% CO2. The isolated cells were placed in DMEM containing 10% fetal bovine serum (FBS), 100 units/ml penicillin, 100 μg/ml streptomycin, at 37 °C with 5% CO2. Only passages 1-3 of the obtained TDSPCs were used in this study. Results of at least three independent experiments are presented.

#### Tendon-like constructs

In order to better mimick the tendon’s natural environment, we performed most of our experiments using 3D-collagen embedded tendon cell cultures. These artificial tendon-like constructs were established as described by our group (Gehwolf et al., 2019). In brief, 2.5 x 10^5^ rat Achilles tendon-derived cells (passage 2) were mixed with collagen type I (PureCol™ EZ Gel solution, # 5074, Sigma-Aldrich, Vienna, Austria; endconcentration 2mg/ml) and spread between two silk sutures pinned with insect pins in rows on SYLGARD 184 (Sigma-Aldrich) coated petri dishes. To improve formation of the constructs, Aprotinin, Ascorbic acid, and L-Proline were added to the cell culture medium. After contraction of constructs over the course of 11 days, 10ng/ml IL-1β (PeproTech, Vienna, Austria), 10ng/ml TNF α (Invitrogen, Carlsbad, USA) and a combination of both cytokines, respectively, was added to the culture medium. After incubation for 24 hours constructs were harvested and stored either in TRIReagent (Sigma-Aldrich, Austria) for further qPCR analysis, fixed in 4% paraformaldehyde for immunohistochemical analysis or frozen at −80°C for subsequent western blot analysis.

#### Animals

C57BL/6 mice (males, 10-12 weeks old, 20-25g) were purchased from the Charles River Laboratories. All animals were acclimatized to standard laboratory conditions (14-h light, 10-h dark cycle) and given free access to rodent chow and water.

Colony-stimulating factor 1 receptor *(Csf-1r)-GFP* and C-X3-C motif chemokine receptor 1 *(Cx3cr1)-GFP* transgenic mice were kindly provided by Dr. Stella Autenrieth from the Medical Clinic of the University of Tübingen and by Prof. Thomas Langmann from the Eye Clinic of the University of Cologne.

Female, 12 week old Fisher rats were purchased from Janvier Labs (France, Europe).

#### Human tendon tissue

Human Semitendinosus tendons available in the course of cruciate ligament reconstructions were provided by the local university clinic after an Ethics approval (E-Nr. 2374) by the local government and prior patients’ informed consent.

#### Preparation of tissue sections

Mouse Achilles, human semitendinosus tendons and rat tendon-like constructs were fixed in 4% paraformaldehyde for 12 hours at 4 °C, and after several washes in phosphate-buffered saline (PBS) and cryo-preservation in 30% sucrose in PBS embedded in cryomedium (Surgipath Cryogel^®^, Leica Microsystems, Vienna, Austria). Subsequently, 12 μm cryosections were prepared using a Leica CM1950 cryostat.

#### Histology and Immunohistochemistry

For descriptive histology cryosections were stained either using Hematoxylin & Eosin or Alcian Blue stain according to standard protocols. In brief, after staining the sections with Weigert hematoxylin for two minutes, the staining was stopped with 1 % acetic acid including a short differentiation step by shortly dipping the slides into HCl/ethanol. After blueing the sections under running tap water for 10 minutes, sections were stained with 1 % eosin Y solution for 1 minute and again immersed in 1 % acetic acid to stop the staining reaction. Subsequently, the sections were dehydrated in an increasing ethanol series (70%, 96%, 2x 100%) and incubated twice in Rotihistol. Finally, sections were coverslipped with mounting medium.

For Alcian Blue staining, sections were incubated in Alcian Blue solution (pH 2.5) for 15 min, rinsed in tap water and counterstained with neutral red stain for 1 min. Finally, sections were rapidly dehydrated in absolute alcohol, cleared in Roti-Histol (Carl Roth, Karlsruhe, Germany) and mounted in Roti-Histokitt (Carl Roth, Karlsruhe, Germany).

Immunohistochemical detection of immune cell-related markers was performed on cryosections of tendons and tendon-like constructs, respectively. After a 5 min rinse in tris-buffered saline (TBS; Roth, Karlsruhe, Germany) slides were incubated for 1h at room temperature (RT) in TBS containing 10% donkey serum (Sigma-Aldrich, Vienna, Austria), 1% bovine serum albumin (BSA; Sigma-Aldrich, Vienna, Austria), and 0.5% Triton X-100 (Merck, Darmstadt, Germany). Followed by a 5 min rinse, slides were subsequently incubated for double or triple immunohistochemistry (overnight at 4°C) with antibodies directed against FKN/CX3CL1 C-X3-C motif chemokine ligand 1 (CX3CL1, #ab25088, Abcam, Cambridge, UK; 1: 100), CX3C chemokine receptor 1 (CX3CR1, #orb10490, Biorybt, Cambridge, UK; 1:100), Cluster of Differentiation 68 (CD68, #sc20060, Santa Cruz, Dallas, USA; 1:50), Cluster of Differentiation 163 (CD163, #ab182422, Abcam, Cambridge, UK; 1:100), epiregulin (EREG/aa1-162, #LS-C314859, LSBio, Seattle, USA; 1:100; #ab195620, Abcam, Cambridge, UK; 1:100), EGF-like module-containing mucin-like hormone receptor-like 1 (F4/80, MCA497RT, Serotec, Oxford, UK; 1:100) and major histocompatibility complex II (MHCII, #ab157210, Abcam, Cambridge, UK; 1:100), all diluted in TBS, BSA, and Triton X-100. After a rinse in TBS (four times 5 min) binding sites of primary antibodies were visualized by corresponding Alexa488-, Alexa568-, or Alexa647-tagged antisera (1:500; Invitrogen, Karlsruhe, Germany) in TBS, containing 1% BSA and 0.5% Triton X-100 (1h at RT) followed by another rinse in TBS (four times 5 min). Some of the slides received an additional nuclear staining using 4’,6-Diamidino-2-phenylindol dihydrochlorid (DAPI). For that, slides were incubated 10 min (1:4000, stock 1 mg/ml, VWR, Vienna, Austria) followed by a rinse in PBS (three times 5 min). All slides were embedded in Fluoromount™ Aqueous Mounting Medium (Sigma Aldrich, Vienna, Austria). Negative controls were performed by omission of the primary antibodies during incubation and resulted in absence of immunoreactivity.

#### In situ phagocytosis assay

Rat flexor tendons (n=3) were freshly isolated and halved lengthwise by a scalpel. The tendons were placed in a 12 well cell culture dish with the cut surface pointing upwards in Minimum essential medium supplemented with 10 % fetal bovine serum, exposing the tendon proper. pHrodo™ Green S. aureus Bioparticles™ conjugate for Phagocytosis (#P35367, Thermo Fisher Scientific, Massachusetts, USA) were added to the tendons at a final concentration of 100 μg/ml. These particles are non-fluorescent outside the cell at neutral pH, but fluorescent (488nm) at acidic pH such as in phagosomes, thus allowing to identify cells with phagocytic activity.

After 24 h, the tendons were counterstained with DAPI for 5 minutes and analyzed by confocal microscopy.

#### Confocal imaging

Confocal imaging was performed using a LSM1 700 confocal microscope (Zeiss) equipped with 405 nm (5 mW fiber output), 488 nm (10 mW fiber output), 555 nm (10 mW fiber output) and 639 nm (5 mW fiber output) diode lasers, a main dichroic beam splitter URGB and a gradient secondary beam splitter forLSM 700 using a 10x EC Plan-Neofluar (10x/0.3) or a 20x Plan-Apochromat (20x/0.8) objective (Zeiss, Munich, Germany). Image acquisition was done with ZEN 2010 (Zeiss), and image dimensions were 1024×1024 pixels with an image depth of 16 bit. Two times averaging was applied during image acquisition. Laser power and gain were adjusted to avoid saturation of single pixels. All images were taken using identical microscope settings based on the secondary antibody control stainings.

#### Quantitative RT-PCR

Total RNA was isolated from tendon-like constructs (n=5 animals, 2 constructs each) using TRI^®^ Reagent (Sigma-Aldrich; Vienna, Austria) according to the manufacturer’s protocol. RNA yield was quantified using a Nanodrop 2000C (ThermoFisher Scientific, Vienna, Austria) and RNA integrity was verified using an Experion Automated Electrophoresis system (Biorad, Munich, Germany). A minimum requirement of the RNA quality indicator (RQI) >7.5 was chosen.

qRT-PCR was performed as described by Lehner et al. using TaqMan^®^ assays from IDT (Integrated DNA Technologies, Coralville, IA, USA) targeting all genes listed in Table 1 (Lehner et al., 2016). Amplification conditions were 50 °C for 2min, 95 °C for 10min, followed by 40 cycles of 95 °C for 15 s and 60 °C for 1 min. All samples were run in duplicate. CQ values were analyzed using qBasePlus v. 2.4 (Biogazelle NV, Zwijnaarde, Belgium) and normalized relative quantities were calculated by normalizing the data to the expression of previously validated endogenous control genes as described by Vandesompele et al. (Vandesompele et al., 2002). As housekeeping genes eukaryotic translation initiation factor 2B subunit alpha *(Eif2b1),* polymerase (RNA) II (DNA Directed) polypeptide A *(Polr2a),* and tyrosine 3-monooxygenase/tryptophan 5-monooxygenase activation protein zeta *(Ywhaz)* were used. The normalized quantities were then determined for the candidate genes scaled against the expression values determined for the controls to generate fold changes in expression.

**Table 1.**
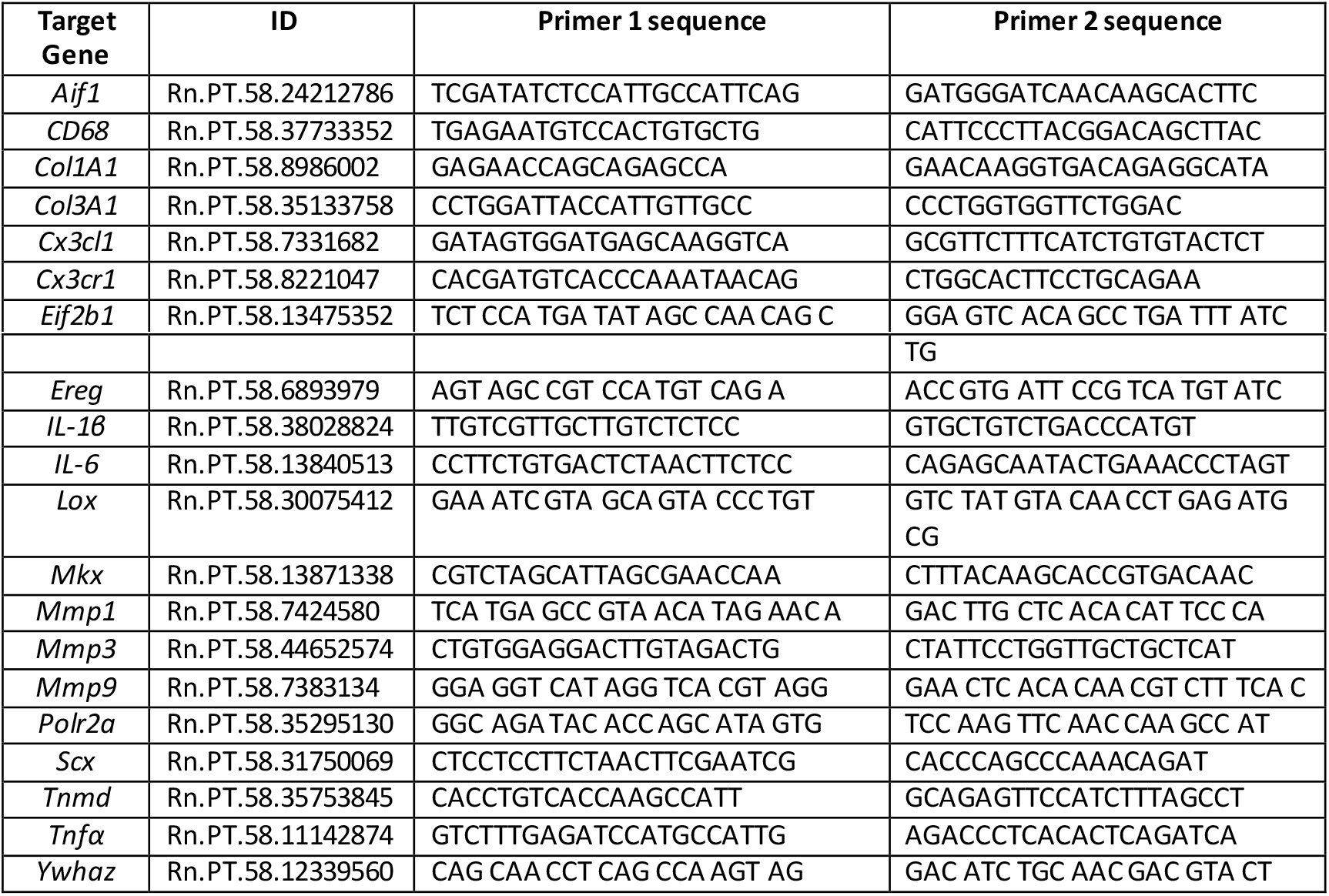

#### Western Blot

Ten to 15 μg of total protein of the tendon-like constructs’ lysate were separated on 10–12% SDS-polyacrylamide gels in Laemmli buffer. Proteins were then transferred to a PVDF membrane (Biorad, Munich, Germany) using 15.6 mM Tris base, 120 mM glycine, and 20% methanol for 1.5 h at 90 V and 4°C. Membranes were blocked in 5% non-fat dry milk powder or 5% BSA hydrolysate in TBS with 0.5% Tween-20, respectively over night at 4°C. Immunodetection was performed using primary antibodies recognizing epiregulin and and CX3CL1 and secondary horseradish peroxidase (HRP)-labelled goat anti-rabbit antibodies, respectively (BioRad, Munich, Germany). Bands were visualized using the Clarity™ Western ECL substrate from BioRad (#170-5060). Band intensities of at least 3 individual experiments were measured densitometrically and normalized to whole protein using the Image Lab Software 5.1 from BioRad (Biorad, Munich, Germany).

#### Migration assay

In order to examine a potential role of fractalkine present in tendon cells on migratory processes, we performed a migration assay using AZD 8797 (Axon Medchem, Groningen, Netherlands), a selective, high-affinity small-molecule inhibitor of CX3CR1. To this end, we seeded rat TDSPCs on both uncoated and collagen coated petri dish. Cells were grown to confluence and serum starved at 1 % serum for 24 hours in order to arrest proliferation. The cell monolayer was then scratched by a sterile 200 μm pipette tip and further cultivated in presence and absence of the inhibitor. After 24 hours, images were taken with a microscope and the distance between the wound margins was measured (Cory, 2011).

#### Staistical analysis

All experiments were repeated at least three times. Statistical analyses were performed using GraphPad Prism v.5.04 (La Jolla, CA, USA). Numerical data is presented as means ± standard deviation. One way analysis of variance (ANOVA) for multiple comparisons and 2-sample t-test for pair-wise comparisons were employed after confirming normal distribution of the data (D’Agostino and Pearson omnibus normality test). Non-parametric statistics were utilised when the above assumption was violated and consequently Kruskal – Wallis test for multiple comparisons or Mann–Whitney test to determine two-tailed p-value samples was carried out. Statistical significance was set at α = 0.05.

## Acknowledgements

We would like to acknowledge Dr. Stella Autenrieth from the Medical Clinic of the University of Tübingen, Germany for providing the CX3CR1-GFP transgenic mouse strain and Prof. Thomas Langmann from the Eye Clinic of the University of Cologne for providing the MacGreen (Cfs1r-EGFP) transgenic mice.

## Author contributions

CL, HT and AT designed the research. CL, GS, HT, AW and NW performed experiments. CD, KE and FW provided human biopsy samples. CL, HT, RG and AT drafted and/or wrote the manuscript. CL, HT and AT provided funding. CL, HT, and AT supervised the work.

## Supplement 1

**Suppl. Fig. 1:**

Longitudinal cryo-sections of Achilles tendons from *Cx3cr1-GFP* transgenic mice co-stained with an antibody directed against the Cx3cr1 protein shows a high degree of overlap (merge), confirming the expression pattern of the Cx3cr1-GFP protein.

**Suppl. Fig. 2:**

Doublelabelling of longitundinal cryo-sections of Achilles tendons from *Cx3cr1-GFP* transgenic mice with antibodies directed against the macrophage-related markers CD68 and CD163 revealed coexpression of these markers with the Fkn receptor (see arrows).

**Suppl. Fig. 3:**

Cross sections of an intact human semitendinosus tendon (♂, 22 years) demonstrating the expression of CX3CL1 and epiregulin in the perivascular region (see white arrows) and the tendon proper (yellow arrows).

